# Persistent Firing Neurons in the Medial Septum Drive Arousal and Locomotion

**DOI:** 10.1101/2021.04.23.441122

**Authors:** Endre Levente Marosi, Karolina Korvasova, Felix Ludwig, Hiroshi Kaneko, Liudmila Sosulina, Tom Tetzlaff, Stefan Remy, Sanja Mikulovic

**Author notes:** Equal contribution (Endre Levente Marosi, Karolina Korvasova).

## Abstract

The medial septum and diagonal band of Broca (MSDB) serve as a central hub in an ascending brainstem pathway that conveys sensory and motor signals to the limbic system. However, the cellular and circuit mechanisms underlying these functions remain unclear. Here, we show that transient optogenetic activation of MSDB VGluT2⁺ neurons initiates a structured arousal sequence - beginning with facial movements, followed by pupil dilation and locomotion. Neuropixels recordings reveal persistent MSDB neuronal activity that strongly correlates with arousal-related behaviors. We demonstrate that persistent firing (PF) is an intrinsic property of a subset of MSDB neurons, independent of ongoing synaptic input. PF neurons and putative GABAergic theta-bursting neurons predicted movement initiation, with population activity scaling with initiation magnitude, unlike other MSDB populations. These findings identify PF in the MSDB as a central neural mechanism that orchestrates the transition from preparatory movements to full behavioral engagement, bridging sensory input with locomotor arousal and supporting state transitions within the limbic system.

## Introduction

The brain’s ability to initiate motivational transitions between behavioral states—such as the shift from rest to locomotion—is fundamental to adaptive behavior and survival. These transitions are accompanied by a range of arousal-related behaviors. In rodents, for example, whisking—a tactile sensory behavior—emerges several milliseconds before locomotion and correlates with locomotor speed^1^. Locomotion is also associated with pupil dilation, which enhances visual inputs^2^. The MSDB functions as a critical hub integrating sensory information across cortical and subcortical networks^3^. Recent studies have shown that MSDB neurons display firing patterns predictive of individual steps within the running cycle^4,5^. Furthermore, a causal role in locomotor initiation has been attributed to MSDB neurons expressing the vesicular glutamate transporter 2 (VGluT2)^6–11^. Beyond locomotion, VGluT2 neurons contribute to spatial goal-directed memory^12^, navigation^7^, and regulate wakefulness and exploratory behaviors through projections to the lateral hypothalamus (LH)^13^ and the ventral tegmental area (VTA)^8^. Sustained optogenetic activation of these neurons at theta frequencies effectively entrains the hippocampal theta rhythm, induces and prolongs locomotion beyond the stimulus duration, and affects animal speed in a frequency-dependent manner^6,8^. In addition, VGluT2 neuron activity increases several hundred milliseconds prior to the onset of voluntary running^6^, pointing to a preparatory phase of locomotor initiation. It remains however unclear whether this neuronal activation is specific to arousal-related behaviors, such as whisking and pupil dilation. The fast-timescale activity dynamics of VGluT2 neurons and other MSDB cell types—likely critical for sensory integration and voluntary movement—remain undefined. Although in vitro studies have revealed complex connectivity among MSDB neuronal populations^14^, how VGluT2 neuron activation shapes activity across MSDB circuits in vivo, and which cellular mechanisms drive locomotion-related behaviors, is unknown.

Persistent firing (PF) refers to sustained neural activity that endures for several seconds after stimulus termination, functioning as a neural toggle switch^15^. Its underlying mechanisms have been extensively studied in cortical regions, where PF contributes to plasticity and learning^16^, including working memory^17^ and head-direction system formation^18^. More recently, PF has been implicated in broader processes such as defensive behaviors^19^ and the transformation of sensory input into motor commands^20^. A unifying feature across these functions is the sustained activity of specific neural populations that supports prolonged behavioral output. Given these diverse roles, a critical question for MSDB is whether PF constitutes a cellular mechanism underlying the transition from rest to locomotion—a process expected to depend on sustained neuronal activation across the second-long behavioral sequence linking arousal to movement onset. Brief whisking bouts and pupil microdilations are well-established markers of micro-arousal episodes, reflecting transient attentional shifts that punctuate quiescent states. Whether MSDB activity plays a causal role in the in the initiation and maintenance of such micro-arousal events, however, remains unresolved.

Here, we show that brief activation of MSDB VGluT2 neurons triggers a well-defined sequence of behavioral and physiological events, beginning with facial movements and followed by pupil dilation and locomotion. Neuropixel recordings revealed a subpopulation of PF neurons whose activity reliably predicted both facial movement and locomotor onset, implicating them in the coupling of arousal and movement initiation. Remarkably, PF neurons displayed intrinsic firing properties independent of synaptic input, as confirmed by pharmacological experiments in acute brain slices. In addition, distinct recruitment patterns of MSDB units—particularly PF neurons and putative GABAergic theta-bursting neurons—predicted the success of locomotor initiation. Collectively, these findings demonstrate that dynamic interactions among MSDB neuronal subtypes are behaviorally relevant, and they establish persistent activity as a key mechanism linking arousal to locomotion-related behavioral responses.

## Results

### Photoexcitation of MSDB VGluT2 neurons elicits facial movement, pupil dilation and locomotion

Previous studies have shown that optogenetic activation of MSDB VGluT2 neurons induces run initiation after prolonged stimulation (>10 seconds) at theta frequency^6–8^. We therefore first investigated whether a shorter, physiologically relevant activation paradigm, applied in the absence of theta-modulated stimulation, would likewise influence animal behavior. In order to test how the length of the stimulus affects locomotion induction in head-fixed mice running on a treadmill, we injected AAV1-EF1a-DIO-ChR-EYFP or an appropriate control construct (AAV1-EF1a-DIO-EYFP) into the MSDB of VGluT2-cre mice, and implanted an optical fiber to deliver 473 nm laser light. In parallel, we investigated the effect of this activation on other locomotion-related modalities, including the face movement (FM) and pupil change (PUP) using Camera Motion Energy analysis and pupillometry, respectively (Fig. 1a, see Methods for details). Two types of stimulus protocols were applied: a previously-used 10s long stimulation at 8 Hz theta frequency^6,8^ or 1s continuous wave (cw) pulses with 2 min inter-stimulation period (Fig. 1b). Compared to control animals, prolonged 8 Hz stimulation induced locomotion with a significantly higher success rate, a trend toward longer duration, and comparable latency. In contrast, transient 1s stimulation produced a success rate indistinguishable from chance and did not affect locomotion duration (Fig. 1c-g). Animals exhibit preparatory behaviors, such as facial movements, prior to locomotion initiation^1^, potentially related to exploration^8^. Additionally, pupil changes are observed during natural running^21^. Therefore, we tested whether the 1s stimulation protocol influenced these behaviors as well. Unlike running, both facial movement and pupil responses were reliably elicited by 1s MSDB VGluT2 stimulation, even in the absence of locomotion induction (Fig. 1h-j). These responses also exhibited a consistent, behaviorally relevant difference in their latencies (Fig. 1k). However, the durations of these three behaviors did not significantly differ (Fig. 1l), suggesting that VGluT2 neuron activation may simultaneously modulate these behavioral responses. To examine why identical stimulation consistently evoked whisking and pupil responses, but did not invariably trigger locomotion, we compared FM and PUP signals between successful (FMr/PUPr) and unsuccessful (FMnr/PUPnr) run initiation trials (Fig. 1h,i). The absolute peak magnitude of both FM and PUP reliably predicted locomotor initiation, with significantly greater values observed during successful compared to unsuccessful trials (Fig. 1m). Moreover, successful trials were characterized by earlier onset of facial movements (Fig. 1n), and by longer-lasting facial and pupil responses relative to no-run trials (Fig. 1o).

**Figure 1.**
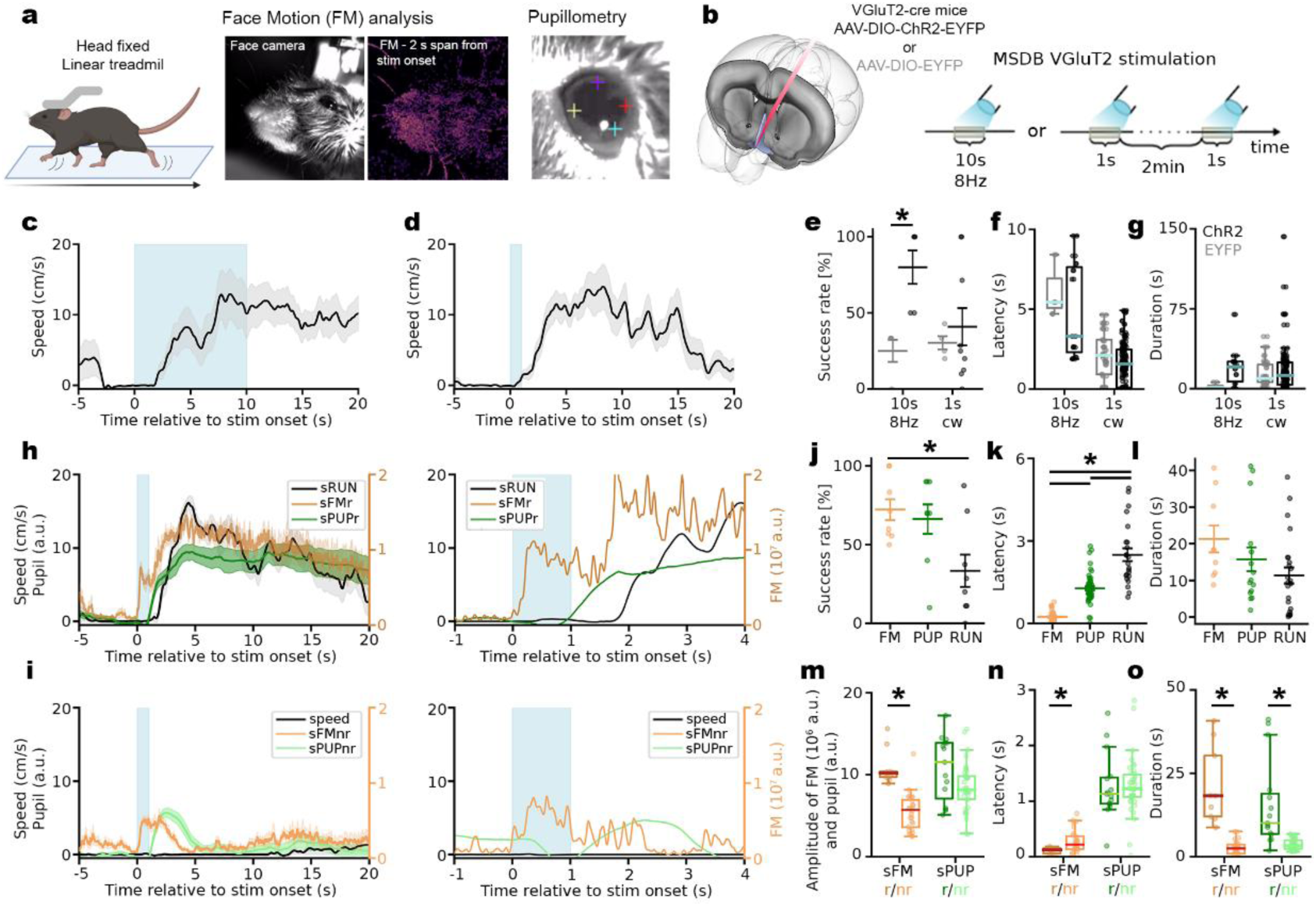
Short, 1 s stimulation of MSDB VGluT2 neurons elicits locomotion, face movement, and pupil dilation. **a** Experimental setup and behavior analysis, (left) running of the animal was tracked on a linear treadmill, (middle) facial movement signal is analyzed from the face camera recordings, example heatmap of 2 s FM shown, (right) pupillometry with DeepLabCut. **b** (left) Placement of the electrode combined optic fiber to the MSDB of ChR2 or control virus injected animals, (right) stimulation protocols. **c** Average speed during and around 10s optical stimulation of VGluT2 neurons at 8 Hz, **d** same as in **c** with 1s continuous wave optical stimulation. **e** Success rate of the stimulation protocols. **f** Latency of run initiation from stimulation onset comparing the protocols and the ChR2 with control. **g** Duration of elicited running. **h** (left) Average trace showing a single-trial successful initiation of stimulated run (sRUN), face movement during stimulated run (sFMr) and pupil dilation during stimulated run (sPUPr) across all animals, and (right) the same data on different time scale. **i** same as in **h**, but for non-successful run initiation (nr). **j-l** success rate, latency and duration of face movement (FM), pupil dilation (PUP) and running speed (RUN). **m-o** Comparison of FM magnitude and pupil amplitude, latency, and duration between successful (darker toned color) and non-successful (lighter toned color) run initiation traces.

Next, we compared voluntary (v) and stimulated (s) behaviors, distinguishing whether they were followed by run initiation or not (sFMr/nr, vFMr/nr, sPUPr/nr, vPUPr/nr), and analyzed their relationships (Supplementary Fig. 1). Locomotion speed and running duration did not differ between voluntary and induced runs (vRUN, sRUN) (Supplementary Fig. 1a,b). However, the latency of preparatory pupil dilation relative to run onset diverged: during voluntary behavior, pupil dilation typically lagged behind run initiation, whereas during stimulation it reliably preceded it (Supplementary Fig. 1c). This indicates that voluntary locomotion involves a natural preparatory period in which pupil dilation is temporally aligned with run onset. Facial movement preceded running in both conditions, but pupil dilation could occur either before or after run initiation (Supplementary Fig. 1c).

The magnitude of preparatory facial movement did not differ between voluntary and stimulated runs; however, stimulation evoked stronger stationary facial movements than voluntary behavior. Similarly, stimulation induced larger pupil dilations than voluntary behavior, both with and without subsequent running (Supplementary Fig. 1d–f). In addition to amplitude, the duration of stimulated versus voluntary facial movements and pupil changes also differed significantly (Supplementary Fig. 1g). Nonetheless, when these behaviors were followed by successful run initiation, their durations converged, consistent with the similarity in sRUN and vRUN durations (Supplementary Fig. 1g). These results indicate that stimulation of the MSDB VGluT2+ population recruits circuits that link whisking, running, and pupil dynamics—three key modalities of locomotor behavior.

Cross-correlation analysis of voluntary behaviors revealed strong coupling between these modalities (Supplementary Fig. 1h). Whisking reliably preceded running, whereas pupil dilation followed both whisker movement and run initiation (Supplementary Fig. 1i,j). Peak cross-correlation lags for RUN × FM and FM × PUP were consistent across animals, but RUN × PUP correlations were more variable, suggesting that whisking acts to synchronize pupil dilation with run onset. Notably, MSDB VGluT2+ stimulation altered this behavioral sequence: stimulation produced the order FM → PUP → RUN (Fig. 1k, Supplementary Fig. 1c), whereas voluntary behavior more often followed FM → RUN → PUP (Supplementary Fig. 1c,i,j). Together, these findings suggest that MSDB VGluT2+ stimulation induces a distinct locomotor state, qualitatively different from voluntary initiation.

### Identification of persistently active neurons in the MSDB

To investigate the neurophysiological correlates of MSDB VGluT2 stimulation–induced behaviors, we performed Neuropixels (NPX) recordings in VGluT2-ChR2 and control animals (Fig. 2, Supplementary Fig. 2). To ensure activation of VGluT2 neurons along the full MSDB axis, we used tapered optical fibers, which distribute light both at the tip and laterally along the fiber shaft. By attaching the optical fiber directly to the NPX probe, we achieved light delivery across the entire electrode shank. Prior to insertion, the NPX shank was coated with the lipophilic dye DiI, enabling post hoc reconstruction of recording sites (Supplementary Fig. 2a). This approach yielded large-scale recordings spanning the MSDB depth, capturing hundreds of units per animal (138 ± 17 units, n = 8; total 1,223 units; Fig. 2a,b). Importantly, this configuration also allowed us to examine network recruitment in relation to locomotor speed, facial movements, and pupil size, and to compare neural dynamics between stimulated and voluntary behaviors (Fig. 2a,b; Supplementary Fig. 2). We hypothesized that the preparatory activation of MSDB neurons reported previously^6,12,22^ might be associated with whisking. To test this, we isolated voluntary stationary facial movement events and found robust MSDB network engagement, with both positively and negatively modulated units at whisking onset (Fig. 2a). Interestingly, these whisking events were consistently followed by pupil microdilations, a hallmark of microarousal and cortical state transitions associated with enhanced information processing^23,24^. Stimulation of MSDB VGluT2+ cells produced longer whisking bouts than voluntary events (Fig. 2a), accompanied by increased unit firing rates (Supplementary Fig. 4a,b). The same units were recruited with similar polarity before run onset during isolated vRUN events (Fig. 2b), and firing rate modulation did not differ between vRUN and sRUN, indicating that running is a secondary consequence of stimulation following whisking (Supplementary Fig. 4c,d). We next characterized the cellular profiles of units engaged during voluntary whisking and running (Supplementary Fig. 2).

**Figure 2.**
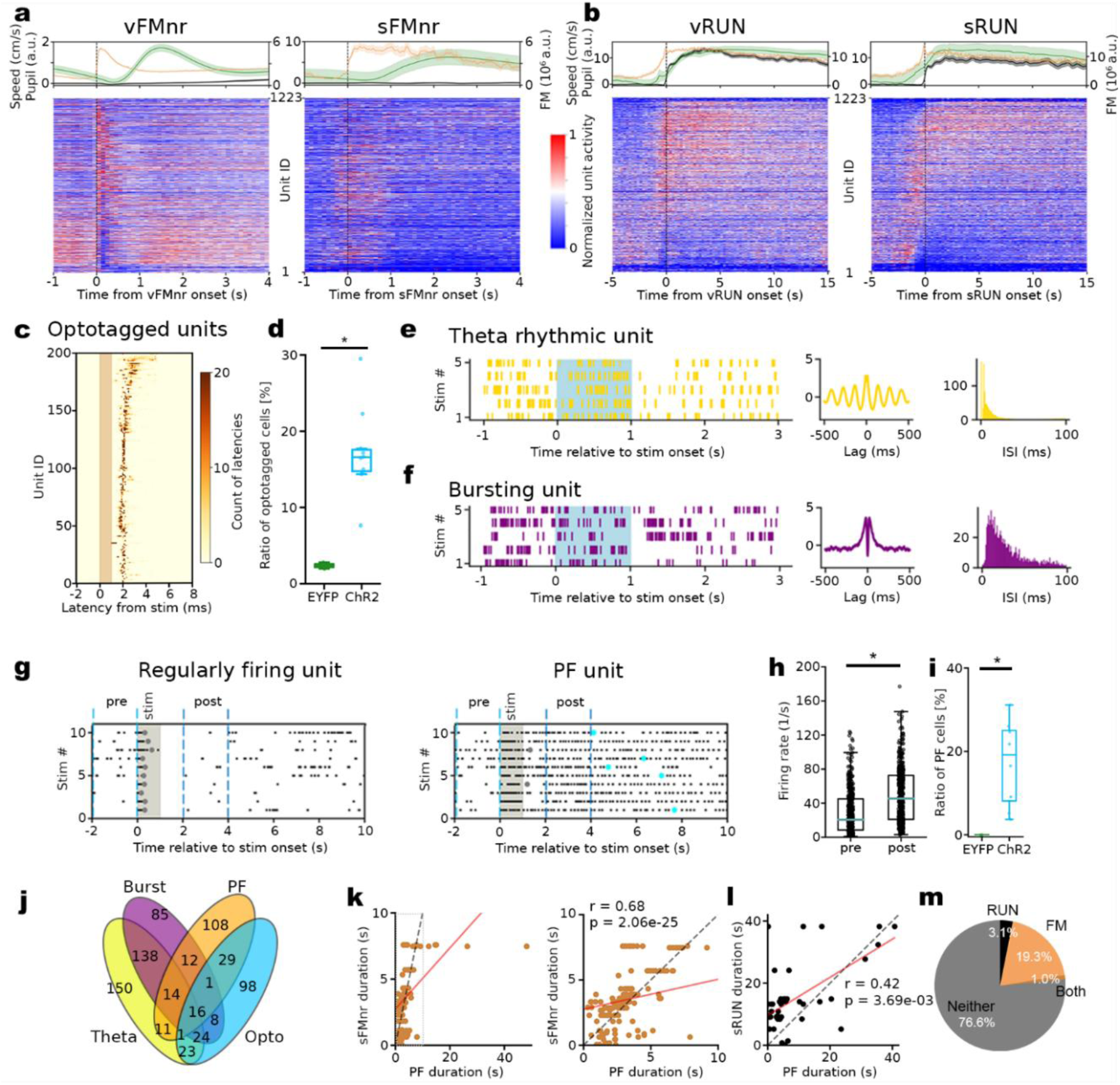
Relationship between behavior and cell-type specific firing in the MSDB. **a** (top, left) Average speed, FM and pupil signals around voluntary stationary whisking (vFMnr) onsets, and (right) around stimulated stationary whisking (sFMnr), (bottom) heatmap of normalized spike rate histogram of the recorded units sorted by rate change before and during FM. **b** Similar to **a**, but with (left) vRUN and (right) sRUN. Unit IDs are conserved between **a** and **b**. **c** Color mapped representation of the response reliability and latency of all optotagged units sorted by mean latency. **d** Ratio of optotagged units. **e** (left) Raster plot of 5 stimulations, (middle) autocorrelogram, (right) ISI histogram of a representative theta rhythmic unit. **f** Similar as in **e**, but with bursting units. **g** Raster plot of 10 stimulation sweeps of a representative unit showing (left) regular firing, and (right) persistent firing. Round markers show the length of elevated firing (gray) shorter or (cyan) longer than 2 s. **h** Comparison of firing rate of PF units before (pre) and 1s after the (post) stimulation. **i** Ratio of PF cells compared to EYFP opsin-free control. **j** Venn-diagram of the cell type of the categorized MSDB units. **k** (left) Correlation of PF length with FM duration, and (right) the same on different x-axis scale with (red) linear fit, and (dashed gray) a diagonal line. **l** Correlation of PF length and run duration. **m** Percentage of PF units showing correlation between PF length and FM and/or run duration.

Using 1-ms light pulses, we first optotagged MSDB VGluT2 neurons by analyzing response latency and spike probability. Units with latencies of 1–5 ms and spike probabilities >10% were classified as optotagged (see Methods; Fig. 2c). Across animals, ∼17% of recorded units were optotagged, consistent with the reported prevalence of VGluT2 neurons in the MSDB^25^ (Fig. 2d). These neurons showed heterogeneous relationships to voluntary whisking and running, including upregulated, downregulated, and non-regulated populations (Supplementary Fig. 3a,b).

Previous studies have reported the existence of theta rhythmic and bursting neurons in MSDB^26,27^. Using a previously established classification approach^28^, we detected both subpopulations (Fig. 2e,f), which showed heterogeneous modulation during voluntary facial movements and running (Supplementary Fig. 3c–f). Notably, optogenetic activation of MSDB VGluT2 neurons revealed a subset of cells that exhibited PF for several seconds after stimulus offset, in contrast to regularly firing units that lacked this phenotype (Fig. 2g). This activity resembles persistent responses reported in cortical and subcortical regions^29,30^. These neurons increased firing rates during both successful and unsuccessful run initiation trials, indicating that their activity was independent of whether stimulation elicited locomotion (Fig. 2h). In ChR2 animals, ∼20% of recorded cells exhibited PF in at least 50% of stimulations (criteria in Methods), whereas no PF cells were detected in controls under our strict classification (Fig. 2i). The PF population consisted predominantly of positively or non-regulated units during stationary facial movements and running, with only a small fraction negatively modulated (Supplementary Fig. 3g,h). For subsequent analyses, we categorized units into inclusive groups with overlapping membership (e.g., optotagged cells could also be classified as PF, theta, and/or burst; Fig. 2j).

Given the robust recruitment of PF cells, we next investigated whether PF duration was correlated with behavioral output. Across all PF neurons, 23.4% were tuned to either facial movement or running (Fig. 2k–m). Specifically, 19.3% showed a strong correlation between firing and FM duration (Spearman’s r = 0.683, p = 2.056 × 10⁻²⁵; Fig. 2k), while 3.1% correlated with run duration (Spearman’s r = 0.420, p = 3.685 × 10⁻³; Fig. 2l). Among all cell classes, PF neurons contained the highest fraction of FM-tuned cells (Supplementary Fig. 3). These results suggest that PF activity supports sustained whisking and/or running triggered by single-pulse stimulation.

### Persistent firing is generated intrinsically in the MSDB

Given the correlation between PF and whisking and/or running, we asked whether PF arises intrinsically within the MSDB or is driven by behavior-related inputs. To isolate the network from behavior, we used acute MSDB slices from VGluT2-Cre mice injected with AAV1-EF1a-DIO-ChR-EYFP (n = 11) or an opsin-free control virus (n = 3), AAV1-EF1a-DIO-EYFP. Slices were incubated in an interface chamber and placed on a perforated 6 × 10 MEA chip (Fig. 3a,b). We applied a stimulation protocol consisting of 1-s continuous light pulses delivered every 2 min, repeated 10 times, with a bath-applied synaptic blocker cocktail introduced midway to assess network contributions (Fig. 3c).

**Figure 3.**
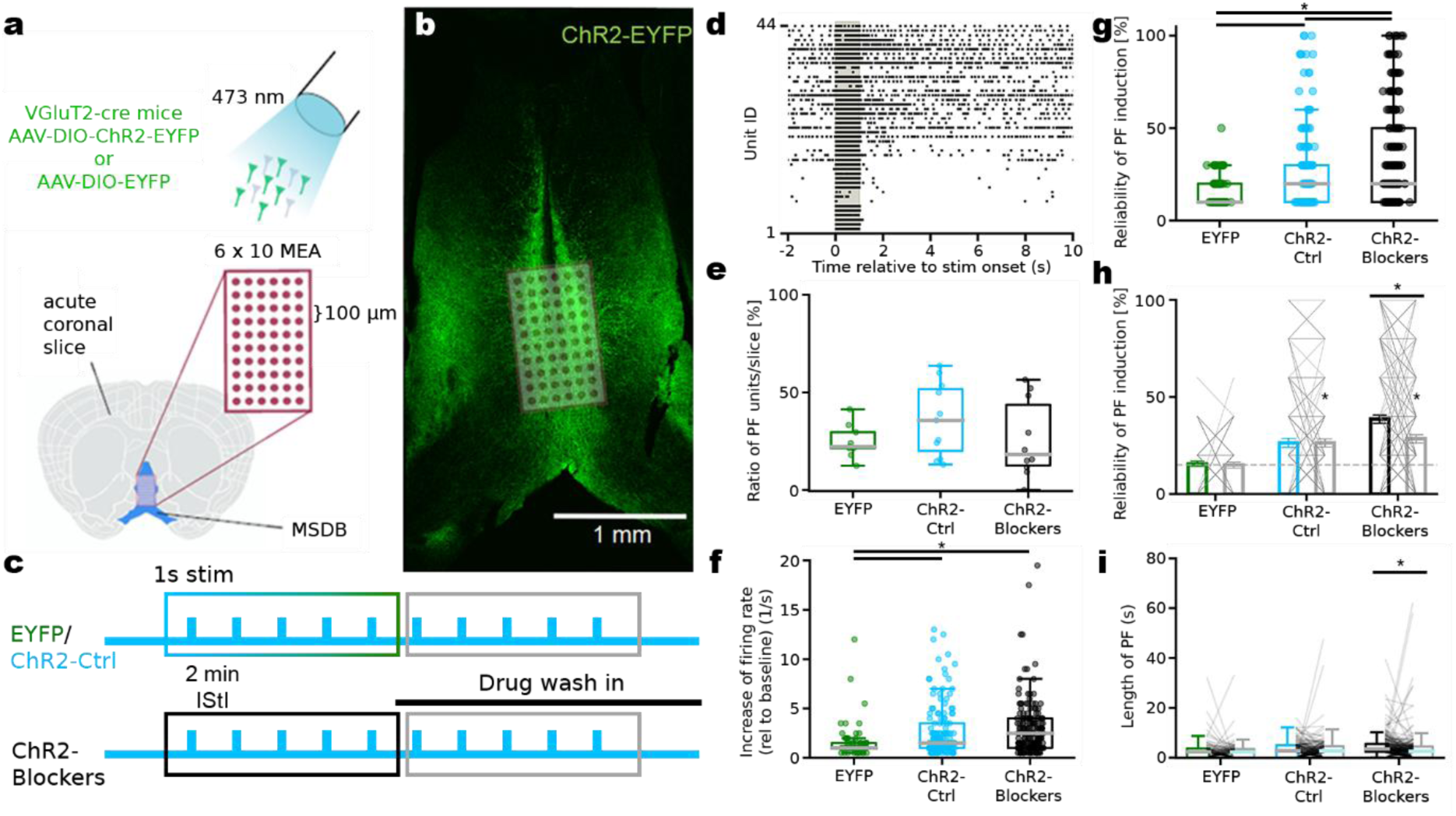
PF is also present *in vitro* in MSDB micronetwork. **a** Experimental design of in vitro slice preparation with Cre-dependent ChR2 expression, 6 x 10 MEA grid. **b** Placement of ChR2 expressing MSDB slice preparation on the MEA grid. **c** Stimulation protocol. **d** Raster plot of recorded units showing regular and normal firing upon photostimulation of a representative experiment. **e** Ratio of detected PF units in recorded slices over different conditions. **f** Photostimulation-induced firing rate change (2 s post-stimulation compared to 2 s pre-stimulation) of PF units were similar between conditions. **g** General reliability of PF elicitation, successfully elicited PF trials over all trials in different conditions. **h** Reliability of PF elicitation, successfully elicited PF trials over all trials in different conditions at the first half (green, cyan, black) and second half (gray) of the experiment. Gray lines show reliability change of individual PF units in the first to second half of the experiment. Dashed line shows the proportion of randomly detected persistent firing by the algorithm in opsin free EYFP slices that was significantly lower than ChR2 expressing slices recorded in ACSF or blocker cocktail. **i** The length of persistent firing decreased with drug application (Blocker cocktail, gray), compared to drug-free baseline (Blocker cocktail, black).

Consistent with our in vivo findings, MSDB neurons increased their firing during stimulation and remained active for several seconds after stimulus offset (Fig. 3d). To test the role of collateral connectivity, we applied a blocker cocktail targeting the major neurotransmitter receptors in the MSDB (mGluR, iGluR, GABAA, GABAB, mAChR, nAChR). This design allowed within-slice comparisons of persistent activity before (stimulations 1–5) and after drug application (stimulations 6–10) (ChR2 + blockers: n = 364 cells, 10 slices, 8 animals). Control conditions included ChR2 slices without drug application (ChR2-ctrl: n = 318 cells, 11 slices, 11 animals) and EYFP slices without opsin expression (n = 217 cells, 12 slices, 3 animals). Persistent activity was observed across all conditions at least once (Fig. 3e). To account for reduced excitability in vitro, we lowered the PF detection threshold from 50% to 10%. Opsin-expressing slices showed stronger post-stimulus increases in firing rate (2–4 s after stimulation) than EYFP controls (Fig. 3f). Across all units, reliability was quantified as the proportion of trials eliciting persistent activity among all trials. PF reliability in ChR2 slices was generally higher than in EYFP control slices, which served as a baseline for detections captured by the algorithm. (Fig. 3g). Importantly, blocker application reduced PF reliability (Fig. 3h) and shortened PF duration (Fig. 3i), yet PF activity persisted, with overall reliability in ChR2 slices remaining significantly above chance relative to control experiments. Synaptic blockers also shortened PF duration (Fig. 3i). Importantly, comparisons across recording conditions were performed within the same slice to control for variability arising from slice preparation. Together, these findings demonstrate that PF can be elicited in MSDB slice preparations in the absence of behavioral output. While local synaptic transmission modulates both PF reliability and duration, persistent activity is not abolished. Thus, intrinsic biophysical properties of MSDB neurons are sufficient to generate PF, with recurrent network dynamics further amplifying and shaping this activity. These findings position the MSDB as a locus capable of intrinsically sustaining activity patterns that may underlie the maintenance of arousal-related states.

### MSDB activity is robustly coupled to facial movements

We next examined how individual MSDB cell types are modulated during voluntary behaviors, specifically stationary whisking and running. Across conditions, all identified cell types exhibited qualitatively similar activity changes; however, the subgroup of non-specific units showed stronger modulation during running compared to stationary whisking (Fig. 4a). When comparing across cell types within the same behavior, burst units, optotagged units, and the pooled population displayed comparable modulation patterns under both conditions. In contrast, theta cells exhibited the strongest negative modulation, with significantly reduced activity during voluntary running relative to the other subgroups, but not during stationary whisking. Conversely, PF cells were predominantly excited during both behaviors, showing modulation patterns that were clearly distinct from all other cell categories (Fig. 4b).

**Figure 4.**
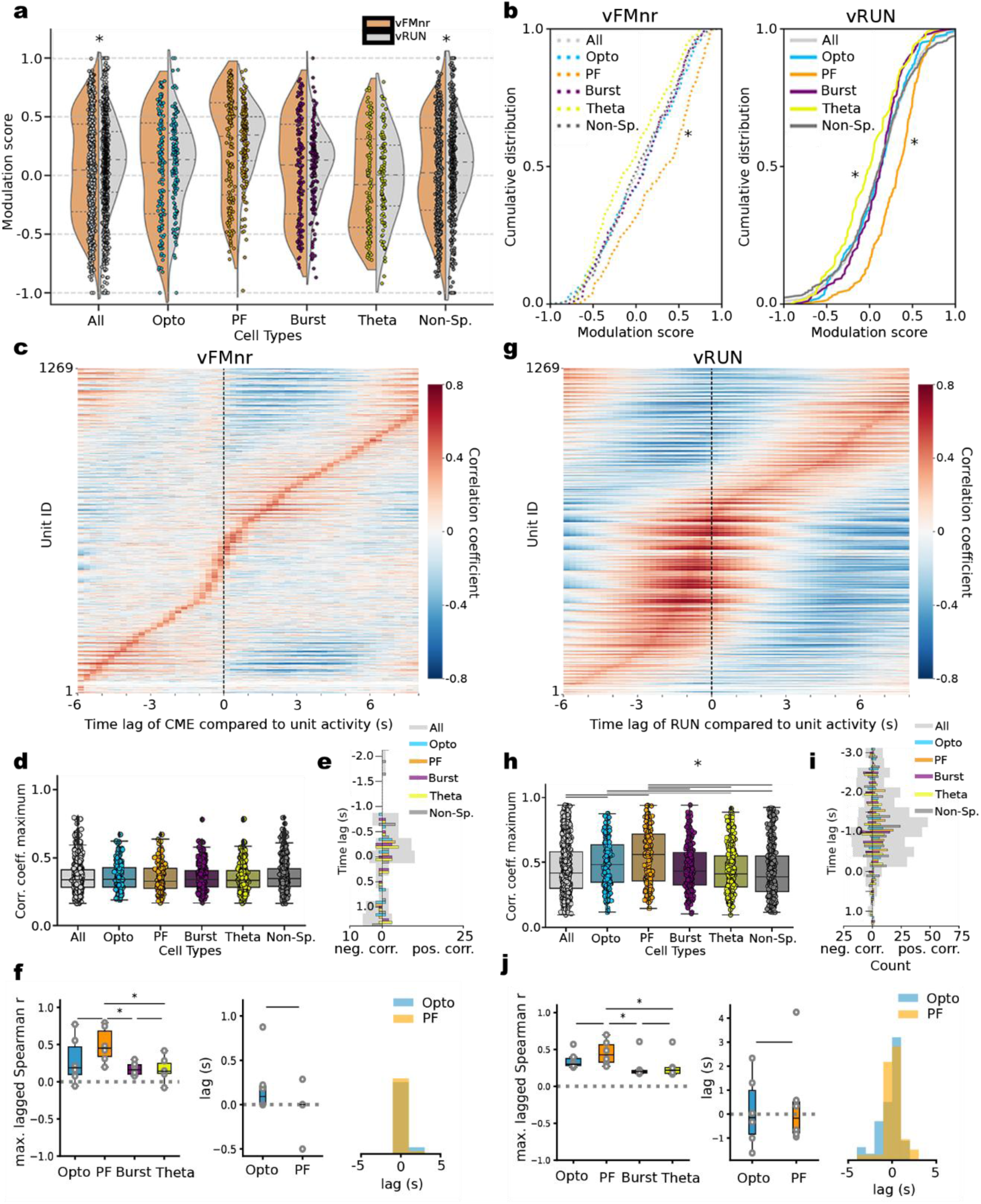
FM is lagged behind MSDB unit activity on a subsecond scale. PF cells are the most involved cell types during voluntary FM and running. **a** Firing rate change representing modulation score distribution plots of all cell types during vFMnr and vRUN with (dashed lines within the distribution plots) median (sparser dashed line in the middle) and interquartile range (denser dashed line on the sides). **b** Cumulative distribution of the modulation scores of identified cell types during (left) vFMnr, and (right) vRUN. **c** Lagged cross-correlation heatmap of activity of all units with vFMnr signal sorted by correlation coefficient maximum. **d** Cumulative barplot of the fractions of FM lag-correlated units of each cell type. **e** Distribution of the lag of maximum positive and negative correlation of all cell types. **f** (left) Maximum lagged Spearman correlation coefficient values of each cell type in 1min windows with 1s bin size, mean per animal (left), the corresponding lags for optotagged and persistent firing cells (middle) and pooled lag values corresponding to Spearman r>0.3 pooled from all animals (right). **g-j** Similar as in **c-f**, but lagged correlation is analyzed between MSDB activity and running.

To further explain why the stimulation has a different effect on FM and RUN behaviors, we investigated the timing precision of neuronal activity using lagged cross-correlation analysis. Visual representation of these cross-correlation values revealed a small subset of MSDB units that were tightly correlated with whisking bouts, with a peak around −250 ms time lag (Fig. 4c). This observation is in line with the previously mentioned stimulation latency of whisking induction (Fig. 1k). On the single unit level, we did not observe any differences in the behaviour of individual unit classes (Fig. 4d,e). However, the population of persistent firing cells strongly synchronizes around the FM onset, unlike the bursting and theta-rhythmic cells (Fig 4f). The synchronization is also visible in the optotagged cell population. On the contrary, majority of the units were recruited during running, and showed correlated activity with this behavior (Fig. 4g-j). On the population level, the activity of persistent firing cells showed moderate correlation with running speed without any major lag, while the bursting and theta rhythmic cell populations remained unaffected (Fig 4j). MSDB activity seems to precede voluntary running with about 1-1.5 s, which value is also representative (Fig. 4i), but not identical to those we measured for stimulated run initiation latencies (Fig. 1k). We also noticed, that almost the entire MSDB network is correlated with running signal, either positively, or negatively, out of which optotagged, confirming the previous findings^6^, and PF cells activity showed the highest degree of correlation with voluntary running (Fig. 4j). Based on the activity profile of MSDB neurons upon self-initiated stationary facial movement events, we identified 5 functionally different subgroups (Supplementary Fig. 5). A subpopulation initiated a peak firing before FM onset (Supplementary Fig. 5a), whereas other groups showed sustained positive or negative modulation matching with the length of elicited FM (Supplementary Fig. 5b,c). A smaller subset of neurons exhibited negative peaks (114/1243 units) (Supplementary Fig. 5d). Interestingly, this group typically displayed theta-bursting activity that abruptly ceased in parallel with FM occurrence. This all suggests that whisking is not only elicited by activation of photosensitive, putatively VGluT2+ cells, but the activity of these, and other cells in the MSDB predicts and modulates whisking behavior.

### Time-lagged coupling between MSDB network activity and pupil dilation

We next analyzed MSDB network activity in relation to pupil dilation. As expected, our stimulation reliably evoked pupil dilation compared to opsin-free control experiments (Supplementary Fig. 6). To disentangle pupil microdilations from locomotion-associated changes, we compared two types of voluntary pupil dilation: run-related and stationary (Fig. 5a). Across conditions, unit activity peaked 1–2 seconds prior to pupil dilation (Fig. 5b–d), consistent with the time lag observed between FM and PUP signals (Supplementary Fig. 1j). A large fraction of units showed significant correlations with pupil dilation (Fig. 5c), with optotagged and PF cells exhibiting stronger correlations than other cell types, while burst cells appeared largely independent (Fig. 5e). Notably, a subset of pupil-correlated cells—similar to the example in Fig. 5a—remained largely inactive during quiescence but fired sparsely (∼1 AP/cycle) in a theta-rhythmic manner during running, tightly aligned with the impending pupil signal. Together, these findings suggest that pupil dilation is a secondary consequence of whisking.

**Figure 5.**
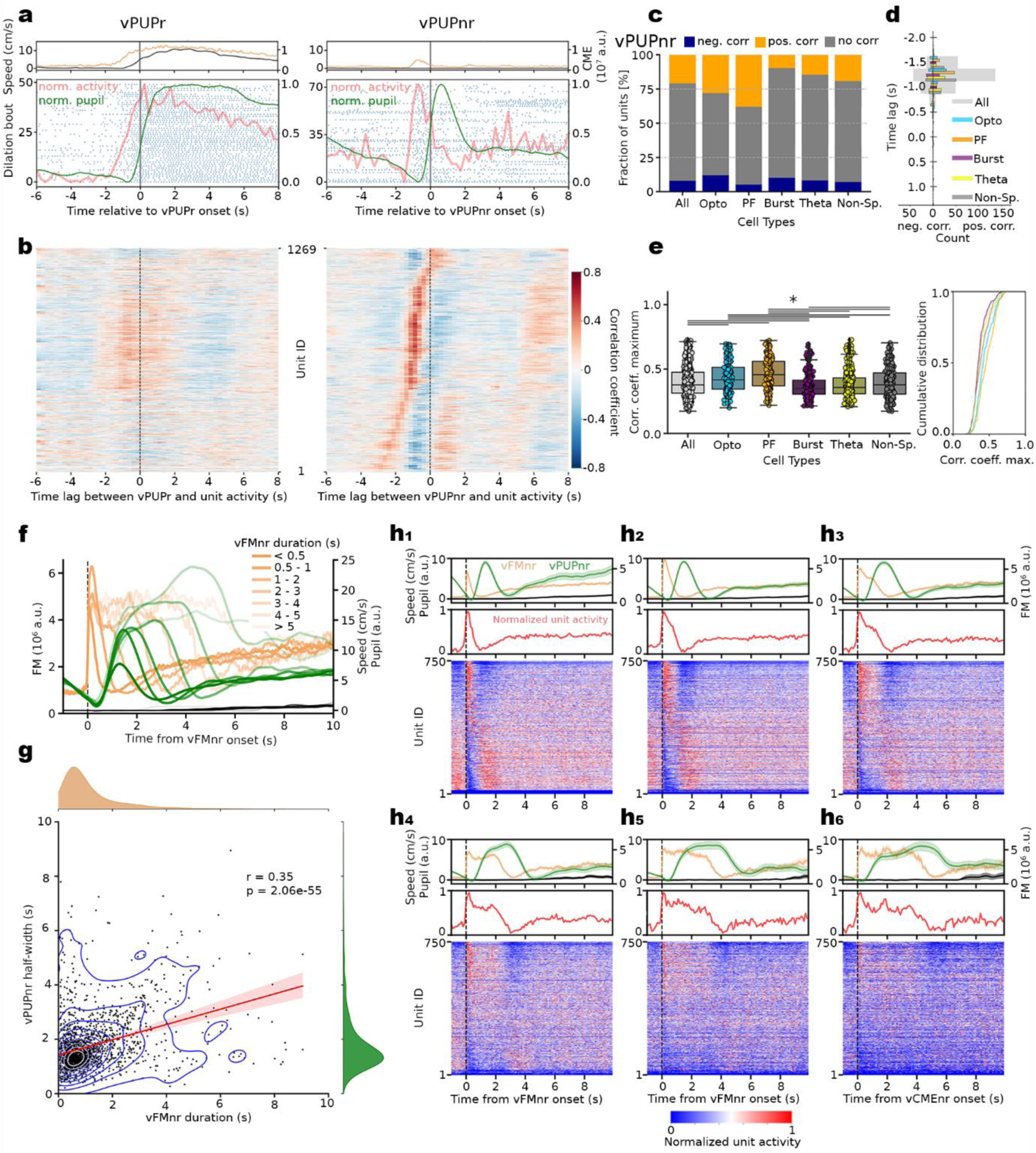
Pupil dilation signal tightly follows FM and MSDB activity. **a** Raster plots, averaged RUN, FM and PUP signals, and average activity histogram of a representative unit centered around (left) vPUPr onset, and (right) vPUPnr onset. **b** Lagged cross-correlation heatmap of all units during to (left) vPUPr onset and to (right) vPUPnr onset sorted by correlation coefficient maximum. **c** Cumulative barplot of the fractions of vPUPnr lag-correlated units of each cell type. **d** Distribution of the lag of maximum positive and negative correlation of all cell types,. **e** (left) Maximum correlation coefficient values of each cell type, and (right) their cumulative distribution. **f** Binned averaged vFMnr and corresponding vPUPnr signal showing linked modulation. **g** Pearson‘s correlation of FM duration and the half-width of the subsequent pupil dilations, with (red) regression line, and (blue) kernel density estimate plot lines. (on the sides) Distribution of the FM (orange), and pupil (green) durations. **h** (top rows) Average speed, FM and PUP signals around binned vFMnr onsets, with (middle rows) averaged activity of the top third (250 units) of the (bottom rows) average normalized activity heatmaps sorted by modulation score. **h1-h6** vFMnr duration bins of 0-0.5 s, 0.5-1 s, 1-2 s, 2-3 s, 3-4 s, 4-5 s bins respectively.

Building on initial observations suggesting a potential relationship between FM and PUP signals (Fig. S1c–j), we next quantified the strength of their coupling. Stationary whisking bouts were binned by duration, and the corresponding signals from all three behaviors were aligned and averaged within each bin. Short bouts of stationary whisking were followed by the smallest and briefest pupil dilations, whereas longer bouts were associated with progressively larger and longer pupil responses (Fig. 5f). Across all stationary whisking events, whisking duration and subsequent pupil dilation duration were significantly correlated (Fig. 5g). Binning cell activity using the same scheme revealed that MSDB population activity closely tracked whisking dynamics. Notably, the average normalized activity of positive sustained units (Supplementary Fig. 5b) closely mirrored the FM signal (Fig. 5h). Together, these findings suggest that pupil dilation is mediated by MSDB network activity, most likely as a secondary consequence of whisking, with the two signals diverging at downstream processing stages.

### General linear model reveals dominant representation of facial movement in MSDB activity

We next trained a general linear model (GLM) to predict neuronal firing from behavioral variables and compared its performance to a conservative conjunctive coding analysis. The conjunctive approach quantified the percentage of run-correlated cells that were also correlated with FM or PUP signals, alongside permutation controls. In the GLM, run initiation episodes were included, and all three behaviors were subsampled to match the number of bins corresponding to neuronal firing rate bins. The model was trained to predict spiking activity from these behavioral inputs (Fig. 6a). Consistent with experimental data, FM alone was the strongest predictor of neuronal firing when behavioral variables were considered individually. However, combining all three behaviors yielded the highest fraction of correctly predicted cells, surpassing FM alone (Fig. 6b). The conjunctive coding analysis further showed that most run-modulated cells were also engaged in FM coding, whereas the reverse relationship was weaker. Specifically, the majority of run-correlated units were co-correlated with FM, with only a smaller fraction co-correlated with PUP (Fig. 6c, left). Conversely, among FM-modulated units, most were co-correlated with run, while only a minority were co-correlated with pupil dilation (Fig. 6c, middle). Finally, the small subset of positively pupil-tuned cells was often co-tuned with run, whereas a relatively larger fraction of negatively pupil-modulated units was also negatively co-tuned with FM (Fig. 6c, right). No differences were observed in the proportions of negatively conjunctive coding neurons across conditions.

**Figure 6.**
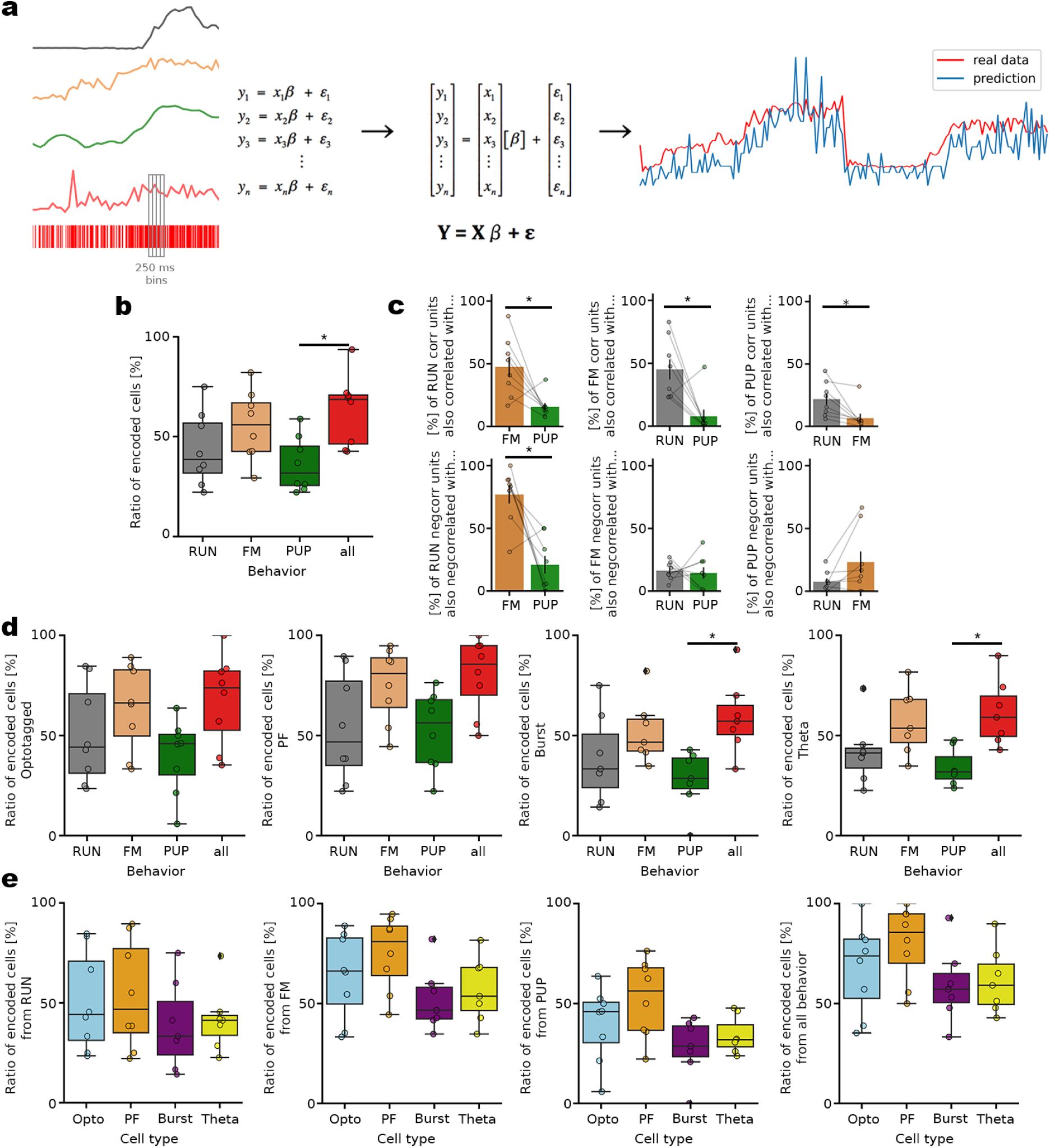
GLM predicts the firing of units from FM with the highest ratio. **a** (left) Input of the GLM (black, RUN; orange, FM; green, PUP; red, unit firing), (middle) schematic of GLM, (right) prediction of the firing of a representative neuron using all the 3 behavioral variables. **b** Ratio of significantly encoded cells by each behavioral signal and all 3 combined. **c** (top) Conjunctive coding of positively (left) speed, (middle) FM, (right) pupil modulated units with the other 2 behavior, (bottom) similarly with negatively modulated units. 100 % is always the total number of modulated units by the reference behavior. **d** Similar to **b**, but for individual cell types (from left to right: optotagged units, PF units, Burst units, Theta units). **e** Fraction of encoded cells of a given cell type by the behaviors (from left to right: encoding from RUN, FM, PUP signals, or all the 3 signals combined).

Notably, PF cell firing was predicted with the highest accuracy from FM (Fig. 6d,e). The GLM analysis thus reinforces our findings, indicating that PF units are the cell type most strongly coupled to behavior, and that among all variables, FM is most prominently represented in the MSDB network.

### Run initiation success predicted by PF and theta-bursting cell activity

To examine the role of the MSDB network in 1-s stimulation–evoked run initiation, we compared successful (run) and unsuccessful (no-run) trials. Among all cell types, PF units showed the clearest difference, exhibiting more sustained activation when stimulation was followed by run initiation (Fig. 7a,b). Given the heterogeneity of PF cells, we hypothesized that recruitment of putatively excitatory PF units (optotagged and unclassified, likely cholinergic) could shift the excitatory–inhibitory (E/I) balance within the MSDB. To test this, we subtracted normalized firing-rate histograms of no-run trials from those of run trials and quantified the difference using the area under the curve, yielding a success score (positive values indicating greater activity during runs, negative values during no-runs). We then focused on the top and bottom 5% of scores. Analysis of theta-bursting cells (putative GABAergic) revealed distinct recruitment patterns: the bottom 5% were preferentially active during no-run trials and suppressed during run trials (Fig. 7c–e), consistent with a feedback mechanism of inhibitory control. Together, these findings suggest that successful run initiation depends on primary recruitment of PF units and secondary modulation by theta-bursting cells, reflecting a shift in E/I balance that facilitates the transition from quiescence to locomotion.

**Figure 7.**
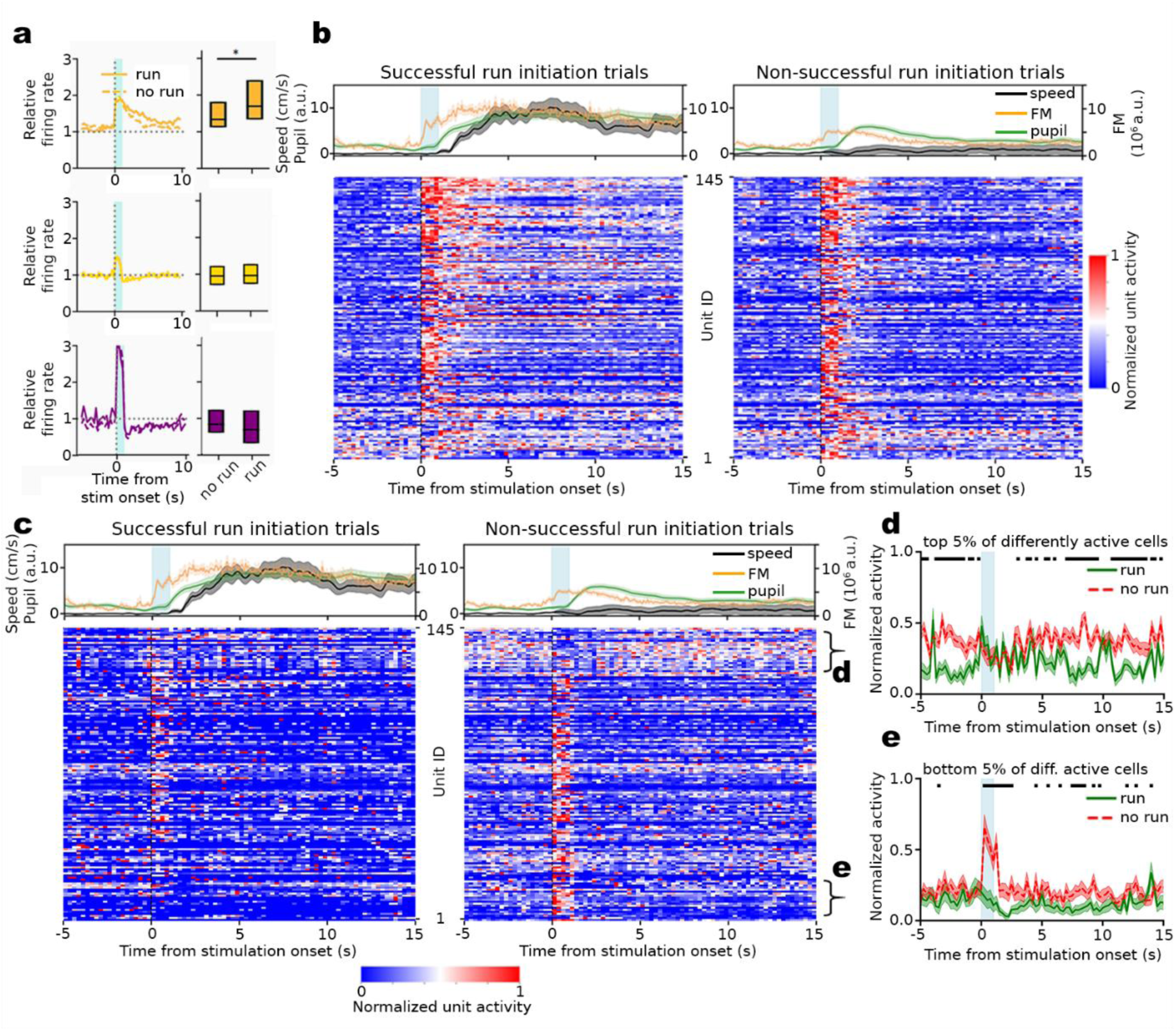
PF and theta-bursting unit recruitment is distinguishable between successful and non-successful run initiation. **a** (left) Firing rate histogram around the 1 s stimulus of (top) PF, (middle) Theta, and (bottom) Burst units, upon successful (solid line), and non-successful (dashed line) run initiations. **b** (top) Average run, FM and PUP signals around stimulation onset of (left) successful, and (right) non-successful run initiations, and (bottom) the subsequent averaged, normalized neural activity of PF units sorted by modulation score. **c** Similar to **b**, but with normalized neural activity of theta-bursting units sorted by success score. Brackets on the right represent the top and bottom 5% of differently active (success-dependent) units. **d** Averaged activity histograms of the top 5% of those units presented on **c** around stimulation when run initiation was successful (green), or not (red). Black squares on the top shows where the activity histograms are significantly different. **e** Similar to **d**, but presenting the bottom 5% of differently modulated theta-burst units.

## Discussion

Previous studies have shown that prolonged stimulation of MSDB VGluT2+ neurons, particularly at theta frequencies, can initiate locomotion^6,8,28^. Here, we explored the effects of shorter, more physiologically relevant stimulation durations. A 1-second stimulation of MSDB VGluT2+ neurons often initiated locomotion, but even failed attempts consistently evoked preparatory behaviors such as whisking and pupil dilation. This suggests that while excitatory–inhibitory balance in the MSDB may be insufficient for running, brief activation reliably recruits arousal-related behaviors. Consistently, MSDB neurons were strongly engaged during self-initiated whisking and pupil dilation, with activity ramping up before locomotion and persisting throughout running (Fig. 2a,b). This activity appeared to entrain whisking as a preparatory signal, potentially coordinating cortical and hippocampal engagement. GLM analysis confirmed whisking as the strongest predictor of MSDB activity (Fig. 6), supporting a role in synchronizing pupil dilation with locomotor onset.

During locomotion, noradrenaline signals precede acetylcholine, highlighting the LC–MSDB pathway in locomotor initiation^23^. Ascending MSDB cholinergic signals further modulate cortical states and enhances visual responses^31^. We found that activation of MSDB VGluT2⁺ neurons induces whisking and pupil dilation even in the absence of locomotion, consistent with microarousal events reflecting cortical state changes measurable through pupil dynamics^32,33^. These state shifts may be the origin of the previously reported ultraslow entorhinal oscillations^34^, which have been implicated in coordinating neural activity across extended timescales and in supporting sequence formation during navigation and episodic memory.

Our results extend prior findings that sounds evoke distinct facial movements^35^ and that aversive stimuli like airpuff^10,28^ or high-decibel noises^36^ evoke MSDB responses, by showing that whisking itself is represented in the MSDB. Selective activation of VGluT2^+^ MSDB neurons induced whisking, suggesting that the MSDB transmits a surprise or arousal signal. As a gatekeeper of arousal and learning, the MSDB’s efferents likely shape distinct behavioral outcomes. MSDB projections to the VTA, LH, lateral and medial habenula, and medial preoptic area, have distinct functions^8,10,36^, indicating segregated circuits that differentially couple locomotion, motivation^9^ and cognitive processes^37^. Notably, aversive stimuli such as airpuff or loud noise rapidly and persistently activate GABAergic and cholinergic MSDB subpopulations^10,11,28^. These findings suggest that arousal states, even without locomotion, may modulate perception and learning^38^. Future work should investigate how MSDB-driven arousal influences learning processes.

Characterization of MSDB neurons revealed a persistently active population whose activity, lasting seconds, was tightly coupled to whisking. This aligns with the time course of PF, suggesting sensory input reaches the MSDB via PF mechanisms. The same neurons were also engaged during voluntary whisking, supporting a physiological role of MSDB circuits in this behavior. Pupil signals showed a delayed correlation with MSDB activity, consistent with their ∼1 s lag behind LC noradrenergic activation^23^, whereas whisking emerged more rapidly (∼200 ms). We propose that whisking-related signals are relayed to cortical areas such as the barrel cortex, accounting for short latencies and allowing synchronization of whisking with locomotion.

In our recordings, whisking and pupil signals were strongly coupled, while running showed greater variability—stimulated runs followed whisking and pupil dilation, whereas voluntary runs were preceded by whisking with pupil dilation aligning to locomotion. This coordination may enhance visual processing during movement. Notably, a subgroup of neurons produced fast FM signals (∼200 ms) while locomotion initiation lagged by several seconds, suggesting PF recruitment bridges preparatory whisking and running. PF activity was typically shorter than a minute, and within our 30-min protocol we observed stable recruitment, downregulation of GABAergic neurons, and increased FM magnitude during successful run initiation. While long-term persistent activity as seen in other regions (e.g., paraventricular thalamus^39^) remains to be tested, longitudinal tracking of PF neurons could clarify how MSDB network recruitment evolves with experience.

Persistent activity can arise from both intrinsic cellular properties and network dynamics. Transient depolarization is sufficient to induce persistent firing even when synaptic transmission is blocked^40–44^, often studied under carbachol to mimic elevated ACh. Other work emphasizes the role of recurrent circuitry^45–51^. In our experiments, transient stimulation of MSDB VGluT2+ neurons evoked persistent firing even without synaptic input or carbachol application, indicating intrinsic mechanisms. Notably, however, persistent firing was more reliable and lasted longer in intact circuits, suggesting that network contributions enhance sustained activity.

Biophysically, persistent firing is thought to arise from calcium-dependent second-messenger signaling pathways^44,52–56^, potentially mediated by calcium-activated non-selective cation currents (I_CAN_) involving TRPM and TRPC channels^57,58^. Neuromodulatory influences are also likely to play a critical role; for example, brief activation of the locus coeruleus induces sustained MSDB activity^59^. Defining the intrinsic biophysical mechanisms underlying persistent firing in MSDB neurons, and determining how neuromodulatory inputs shape these dynamics, will be important directions for future work.

In summary, our findings establish persistent firing in the MSDB as a central mechanism that extends beyond its classically described role in episodic memory and associated cortical circuits. Rather than being memory-specific, MSDB persistent firing regulates locomotor state maintenance and may additionally shape reinforcement learning and anticipatory coding. By synchronizing hippocampal and entorhinal theta oscillations with motor rhythms^4,5,28,60–62^ and coupling to whisking- and pupil-linked arousal, these neurons are positioned to coordinate motor planning with spatial and cognitive processing. We propose that persistent firing neurons within MSDB circuits function as a state-dependent control hub linking preparation to action, thereby enabling flexible, goal-directed behavior in dynamic environments.

## Methods

### In vivo experiments

In vivo experiments were performed in adult, 10-14 week old homozygous VGluT2-cre mice (B6J.129S6(FVB)-Slc17a6^tm2(cre)Lowl^/MwarJ) of both sexes, (The Jackson Laboratory, Bar Harbor, ME USA). Mice were group-housed before, and single-housed after the surgery under 12-hour inverted dark and light cycle. All experiments were performed in the dark phase of the cycle with food and water ad libitum. All experimental procedures were approved by the local authorities of Saxony-Anhalt and performed in accordance of the regulations at Leibniz Institute for Neurobiology in agreement with European Committees Council Directive.

### Surgical procedures

Stereotactic surgery was performed under deep isoflurane anesthesia (Vetflurane 1000 mg/g, Virbac, Carros, France) in a stereotactic frame (MA-6N head-holder, GM4 anesthesia mask, Narishige, Tokyo, Japan) under the guidance of stereomicroscope (M80, Leica Microsystems GmbH, Wetzlar, Germany), and mice were heated to 30°C using a heating pad (Rodent Warmer X2, Stoelting Co, Wood Dale, IL, USA) throughout the surgery. After scalp removal the skull was prepared, dried, a checkerboard pattern was carved into the bone, and phosphoric acid gel (37.5 %, Gel Enchant, Kerr, Kloten, Switzerland) was applied to increase the adhesion surface. A thin layer of Primer and Adhesive (Optibond, Kerr, Kloten, Switzerland) were applied to the skull. A small craniotomy was drilled (Q Basic ST drill, Schick GmbH, Schemmerhofen, Germany; H71.104.003 drill bit, Komet Dental - Gebr. Brasseler GmbH & Co. KG, Lemgo, Germany) above the medial septum (+1.0 mm anterior and 0 mm lateral, relative to bregma), and the sinus was carefully pushed away by the 34G beveled needle of a Hamilton syringe (WPI, Sarasota, FL, USA), with its opening pointing to the midline. Channelrhodopsin (pAAV2.1-EF1a-double floxed ChR2-EYFP-WPR (H134R)) or opsin free control virus (pAAV2.1-EF1a-double floxed EYFP-WPR (H134R) (Addgene, Watertown, MA, USA)) of 400 nl was injected into each of two loci of the medial septum (4.5 mm and 4.0 mm ventral, relative to bregma) with a speed of 100 nl/min using motorized manipulators (Mini 23-XR motors, SM-7 remote control pad, SM-10 control box, Luigs & Neumann GmbH, Ratingen, Germany) and micropump (UMP3 pump, Micro4 pump control, WPI, Sarasota, FL, USA). After injection, a 1 mm diameter circle (centered at AP: 1.0 mm, ML: 0.7 mm relative to bregma) was drawn and slightly carved into the bone to mark the craniotomy sight, and small metal bar was placed above the posterior part of the skull for the acute head-fixed experiments (Luigs-Neumann, Ratingen, Germany). To prepare reference and ground electrode outposts, 3 craniotomies were drilled on the occipital bone, and blunt ending golden pin connected silver wires were immersed into each craniotomy epidural above the cerebellum. The skull was covered up with UV-curable dental cement (Gradia Direct Flo, GC Corporation, Tokyo, Japan), except above the marked window, that was covered by Kwik-sil (WPI, Sarasota, FL, USA) until the preparation for the acute experiment. After the surgery, animals were single-housed with a home cage treadmill to ensure their activity during the experiment. Animals were allowed to recover for 2 weeks before the habituation.

### Recording preparation

After the mice were handled and habituated to headfixation, they were trained on the treadmill until they become good runners (meaning that they covered at least 20 m in 10 min). Before the experiment, mice went under isoflurane anesthesia, and the pre marked 1 mm craniotomy and subsequent duratomy was performed. We made sure that the exposed brain tissue is always covered with saline, while a non-operating (dummy) Neuropixels 1.0 probe (IMEC, Leuven, Belgium) was immersed into the tissue to ensure penetration. After this test, the tissue was covered with 1% low melting point agarose (dissolved in saline) (Agarose low EEO, PanReac AppliChem, Darmstadt, Germany) and Kwik-sil, and mice were allowed to rest overnight, or at least 6h before the acute probe immersion.

### Recording

Prior to the head fixation and preparing the animal to run on a treadmill (length: 360 cm, width: 7 cm, color: black, no cue), the Neuropixels 1.0 probe (IMEC, Leuven, Belgium) combined with optical lambda fiber (Optogenix, Lecce, Italy) with 2.5 mm active length was gently painted with DiI (1 mM, Vybrant, Invitrogen, Thermo Fisher Scientific, Waltham, MA, USA) to later recover the site of recording. The combined probe (Fig S2a) was than immersed slowly into the brain (<5µm/s) with 10° angle until reaching the 5500 µm from brain surface. The probe was held for at least 15 min to stabilize the tissue, and to reach sufficient signal-to-noise ratio. The recording consisted of 5 different protocols, where the 473 nm diode laser (LuxX473-80, Omicron-Laserage Laserprodukte GmbH, Rodgau-Dudenhofen, Germany) was controlled by custom written code (Igor Pro 7, WaveMetrics, Portland, USA). We used 1) 1s continuous wave stimulation with 2 min inter-stimulation intervals (IStI), 10 stimulations, 2) 500 ms cw stimulation with 20 s IStI, 5 or 10 stimulations, 3) 100 ms cw stimulation with 20 s IStI, 5 or 10 stimulations, 4) 5 ms cw stimulation with 20 s IStI, 5 or 10 stimulations, and 5) 10 s long stimulation at 8 Hz with 1 ms pulse width. Except for Supplementary Figure 6, only protocol 1 and 5 were analyzed in this study.

Electrophysiological data was acquired through the Neuropixels on board amplifier, transferred by a tether cable, and sampled at 30 kHz through an acquisition module (PXIe-1071, National Instrument, Austin, TX, USA), and by an interface module (MXI-Express x8, PXIe-8381, National Instrument) was further transferred to the acquisition computer (Z4, HP Inc., Palo Alto, CA, USA), controlled by Open Ephys software (v 0.6.4, Open Ephys Inc., Atlanta, GA, USA). One infrared lamp (LIU780A, Thorlabs Inc., Newton, NJ, USA) and two side cameras (acA2040-90umNIR, Basler, Ahrensburg, Germany) were used to monitor the animals face and body, and the face camera signal was further analysed to acquire pupil signal (DeepLabCut 2.3.5^63,64^), and camera motion energy (custom written code, Python), which value was used as facial movement (FM [a.u.]). The speed of the animal was monitored by a rotation sensor built-in the treadmill (Luigs-Neumann, Ratingen, Germany).

### Data analysis

Data analysis was performed using custom-written scripts in Python v3.6.10, with the packages Numpy v1.19.2, Scipy v1.5.2, Neo v0.7.1, Elephant v0.6.2 and Matplotlib v3.3.2. Spike rate histograms were sorted by modulation score, calculated from the spiking activity during behavior or stimulation relative to baseline. Facial movement and running events were isolated by thresholding 2.5*10^6^ and 0.3 cm/s, respectively. Pupil dilation events were defined as time points at which the first derivative of the pupil signal reached or exceeded 6 px/s.

### Spike sorting

After the recordings, we used Kilosort 3 (https://doi.org/10.1101/2023.01.07.52303665) to preprocess and sort the spikes, and subsequently phy software for manual curation (https://github.com/cortex-lab/phy). Good units were selected based on template waveform, refractory period validation, and amplitude distribution throughout the recording. Units were merged and split manually by the guidance of firing rate maps, waveform, anatomical location, and template principal component charts. The spike times for the good units were extracted to python for further analyses with Npyx package^66^.

### Cell type characterization

- Persistently firing units

Persistent activity was defined as significantly increased firing rate in the time interval 1–3 s after the stimulus offset (i.e. 2–4 s after the stimulus onset), compared to baseline calculated over 2 s before the stimulus onset. Only units showing persistent activity at least 50% in response to the 1 s stimulations (5 out of 10 stimulations) were considered persistent firing units in our in vivo experiments. Persistent activity length was calculated by binning the spike trains to 200 ms bins, and 2 s window before the stimulation was statistically tested against the 2 s window from stimulation onset, then this latter 2 s window was shifted with 200 ms, and p-values were continuously collected. When the list of p-values increased above 0.05 (i.e. the unit activity of the pre-stim window showed no difference from the activity in the sliding window), the last AP from that time point was considered as the end of persistent firing.

- Optotagged units

Optotagged units were identified with protocol 5), a train of 1 ms laser pulses at 8 Hz. Units that showed response at least 10% of the stimulation within 5 ms were considered optotagged. The spikes in the 1s stimulation window were neglected to exclude potential artifacts of the stimulation.

- Theta-rhythmics and theta-bursting units

For identifying theta-rhythmic and bursting neurons, we adapted the method from Kocsis et al., (2022)^28^. First, the autocorrelation function was calculated at a 1 ms resolution, smoothened by a 20 ms wide moving window and normalized to have an integral equal to 1. To assess whether the neuron is theta-rhythmic and/or bursting, we calculated two measures: the rhythmicity index (RI) and the burst index (BI).

Theta-rhythmic neurons were characterized by their auto-correlation function where it shows significant modulation in the theta-frequency range. The RI is based on the normalized difference between the amplitude at the highest positive-lag peak and at the lowest positive-lag trough of the autocorrelation function. To calculate the peak value, we first concatenated segments of the autocorrelation function in the lag intervals (110 ms, 150 ms) and (240 ms, 280 ms), where theta peaks are expected, and calculated maximum denoted as ac_peak_. For the trough values, we similarly concatenated the lag intervals centered in between the expected peak position, i.e. (45 ms, 85 ms) and (155 ms, 195 ms), and calculated the minimum ac_trough_. Then the RI is calculated as

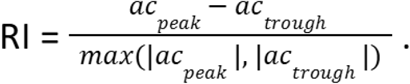

Cells with RI > 0.3 during periods of voluntary locomotion were marked as theta-rhythmic.

For Burst units, we calculated burst indices (BI). Values in the lag interval (15 ms, 40 ms) of the autocorrelation function, denoted as ac_theta_, were compared with values in an extended interval (15 ms, 80 ms) denoted as ac_ext_. The BI was calculated as

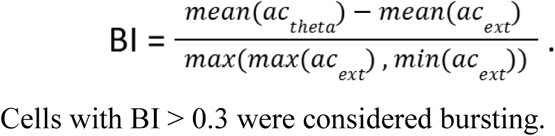

Cells with BI > 0.3 were considered bursting.

### Generalized linear model

To investigate the unit coding of individual or conjunctive behaviors, we used a Generalized Linear Model (GLM) to infer unit activity based on the min-max normalized behavioral variables (FM, speed, pupil diameter). First, we grouped the behavioral variables as well as the spiking data into 20 second long windows centered around each voluntary run initiation, resampled the signals to an equal same sample rate, and finally binned the signals into bins of 250 ms. A GLM for Poisson-distributed observables was then trained on 80% of the time windows for each animal consisting of the behavioral variables as predictors and the binned spiking as outcome variable using the statsmodels Python library (v 0.14.3). We then ran prediction of unit activity based on the remaining 20% of time windows and computed the R^2^ scores between the real and inferred data. To validate if the correlation is significant over chance, we bootstrapped the behavioral data by first dividing the behavioral data into blocks of 19 data points and randomly permuting the blocks, second by circular shifting the time series by a given offset. The procedure was repeated 100 times, and units with the original R^2^ scores higher than 95% of the bootstrapped R^2^ scores were considered as significantly predicted unit activity from the behaviors. Conjunctive coding was annotated to units having significant activity predictions by at least 2 behavioral variables.

## In vitro experiments

### Experimental setup

For MEA recordings of spontaneous and light-induced action potentials, with isofluran, we anesthetized adult (22-48 week old) VGluT2-cre mice that went under virus injection surgery, where similar constructs to in vivo experimental procedures were used. After 26-60 day incubation period, 400 µm thick coronal MSDB slices were cut in ice-cold high sucrose artificial cerebrospinal fluid (ACSF) (mM): 85 NaCl, 75 sucrose, 2.5 KCl, 25 glucose, 1.25 NaH2PO4, 4 MgCl2, 0.5 CaCl2, and 24 NaHCO3; using Leica VT1200S vibratome. After cutting, slices were transferred to an interface chamber (Warner Instruments, Hamden, USA) containing standard ACSF for recovery (mM): 125 NaCl, 3 KCl, 26 NaHCO3, 2.6 CaCl2, 1.3 MgCl2, 1.25 NaH2PO4, and 15 glucose, oxygenated with 95% O2 and 5% CO2. MSDB slices were kept inside the interface chamber on lens cleaning tissue (Grade 105, Whatman, Maidenstone, England) allowing optimal oxygenation due to a laminar flow of preheated (35°C) ACSF above and underneath the slices for at least 30 minutes of incubation until they were used for the experiment when the slices were transferred to the recording chamber for data acquisition.

### Recording

Extracellular waveforms in the MSDB slices in VGluT2-cre mice were recorded with a MEA2100-System (Multi Channel Systems, Reutlingen, Germany, RRID:SCR_014809) on 60pMEA100/30iR-Ti MEAs with round titanium nitride (TiN) electrodes, as described in Sosulina et al. (2021). In detail, the MS slices were positioned onto a 6 x 10 matrix of electrodes, with a spacing of 100 µm and an electrode diameter of 30 µm. ACSF temperature was adjusted to 35°C using heatable PH01 perfusion cannulas together with a TC01 controlling unit (Multi Channel Systems, Reutlingen, Germany). The position of the slice was stabilized by applying a constant negative pressure of the perforated chamber 25 - 30 mBar. Data were acquired with Multi Channel Experimenter (V 2.18.0.21200, Multi Channel Systems, Reutlingen, Germany) at 25 kHz sampling rate with an MEA2100-lite-Interface Board.

### Optogenetic stimulation in MSDB brain slices

Brain slices of VGluT2-cre mice expressing channelrhodopsin in the MSDB were used for optogenetic experiments. Optogenetic stimulation in slices was performed with a light fiber coupled 473 nm diode laser (LuxX473-80, Omicron-Laserage). The light fiber tip was placed at a distance of ≤ 5 mm to the slice. We used a custom-written Igor script stimulation protocol forwarded to the MEA board to synchronize the recording with the stimulation. The 30 min long stimulation protocol consisted of 5 minutes of baseline period before and after the stimulation for sanity control of the condition of the slice, in between 10 pulses of 1s continuous laser stimulation with 2 min IStIs were applied. In pharmacological experiments, ACSF bath solution was changed after the 5th repetition of the stimulus to blocker cocktail containing ACSF solution serving as an inner control for our in vitro experiments (µM): 10 NBQX, 50 D-AP5, 10 SR-95531, 1 CGP52432, 10 Atropin, 0.2 MLA, 1 MCPG. In EYFP experiments the 10 repetition protocol was applied without any change of the standard ACSF bath solution.

### Data analysis

After data collection, slices were transferred overnight to 4% PFA solution and were mounted for confocal tile-scan imaging (SP8, Leica, Germany). Only slices with EYFP expression were then used for data analysis.

### Software

Recordings were exported to HDF5 format and were analyzed with Python. Data were preprocessed using 300-6000 Hz band-pass filter, and common median reference using Spikeinterface package^67^. Spikes were sorted with Tridesclous 2 sorter (https://github.com/tridesclous/tridesclous), spike times combined with stimulation times were further analyzed.

### Statistical testing

In the cases with clearly defined pairs of values (e.g. pre- and post-stimulus value of the mean firing rate), the two-sided Wilcoxon’s signed-rank test was used. In all other cases the non-parametric two-sided Mann-Whitney U-test or Kruskal-Wallis-Test for multiple comparison was applied, as the data were not normally distributed.

## Acknowledgement

We thank Balazs Hangya, Motoharu Yoshida, Petra Mocellin, Jens Kremkow and Janelle Pakan for their valuable comments on an earlier version of the manuscript; Daniela Hill for technical assistance; and all members of the Cognition & Emotion Laboratory for insightful discussions. This work was supported by the LIN Magdeburg Start-Up Package to Sanja Mikulovic and the LIN facility funded through the European Regional Development Fund (ERDF).

## Contributions

E.L.M. performed experiments with inputs from F.L., H.K. and L.S. E.L.M. and K.K. performed data analysis with inputs from T.T., S.R. and S.M. S.M. supervised the project. E.L.M., K.K., and S.M. wrote the manuscript with inputs from all authors.

