## Supplementary figures for "Persistent Firing Neurons in the Medial Septum Drive Arousal and Locomotion"

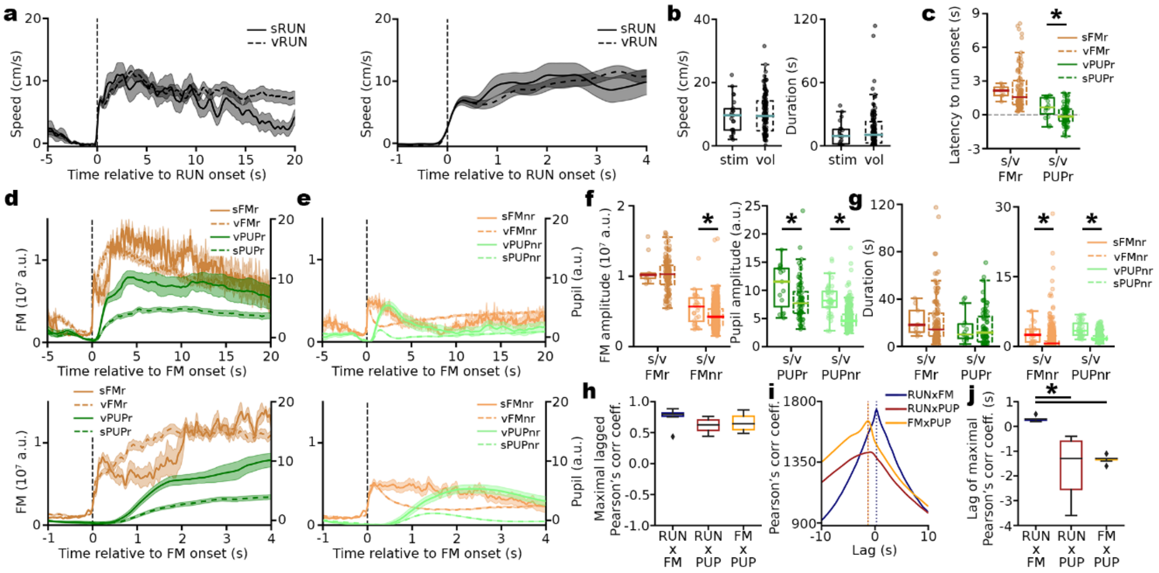


**Supplementary Fig. 1, Comparison of voluntary and stimulated run, FM, and PUP signals. a** Average voluntary and stimulated run traces throughout all experiments. **b** Comparison of speed and duration of voluntary and stimulated running bouts. **c** Latency of whisking and pupil compared to stimulated (solid) and voluntary (dashed) run onset. **d** (top) Average traces of voluntary, stimulated run-preparatory facial movement and pupil signals (vFMr, sFMr, vPUPr, sPUPr) around FM onset, (bottom) the same traces on different time-scale. **e** (top) Average traces of voluntary, stimulated stationary facial movement and pupil signals (vFMnr, sFMnr, vPUPnr, sPUPnr) around FM onset, (bottom) the same traces on different time-scale. **f** (left) FM amplitude of sFMr, vFMr, sFMnr, and vFMnr, (right) pupil amplitude of sPUPr, vPUPr, sPUPnr, and vPUPnr. For legend use **d** and **e**. **g** Similar to **f** but with comparison of duration. **h** Boxplot of maximum of lagged cross-correlation coefficients of the experiments. **i** Average lagged cross-correlation of the 3 behaviors through all experiments. **j** Box plot of the lag of the maximum cross-correlation coefficients of the 3 behaviors compared.

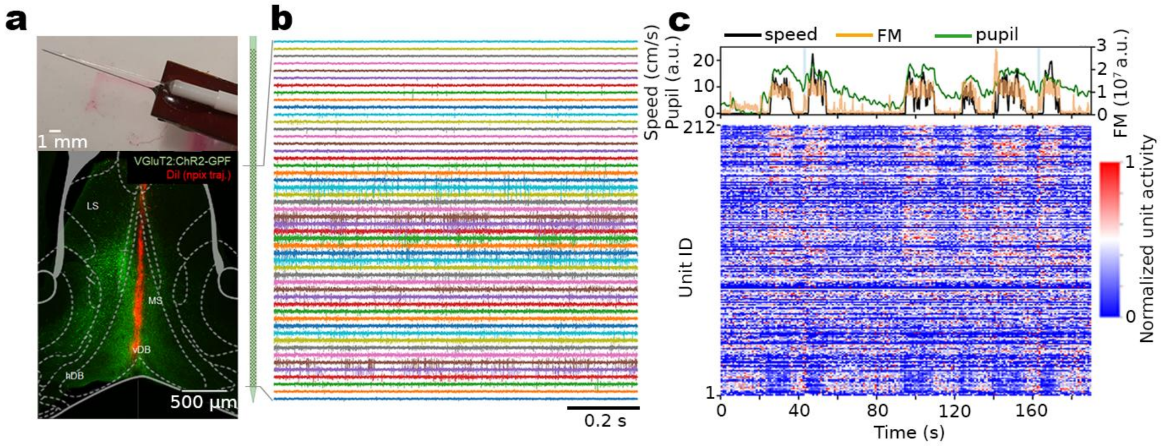


**Supplementary Fig. 2, Design of NPX recordings. a** (top) Self-designed optrode of a tapered optic fiber attached to the NPX probe, (bottom) trajectory of NPX probe of a representative experiment. **b** representative recording cut-out of the shank placed in MSDB. The size of the schematic probe is matched with the histology micrograph. **c** Snippet of a representative experiment with (top) RUN, FM and PUP plotted and (bottom) subsequent normalized neural activity sorted by modulation score calculated before and during running. The light blue lines on the top represent 1s stimulation.

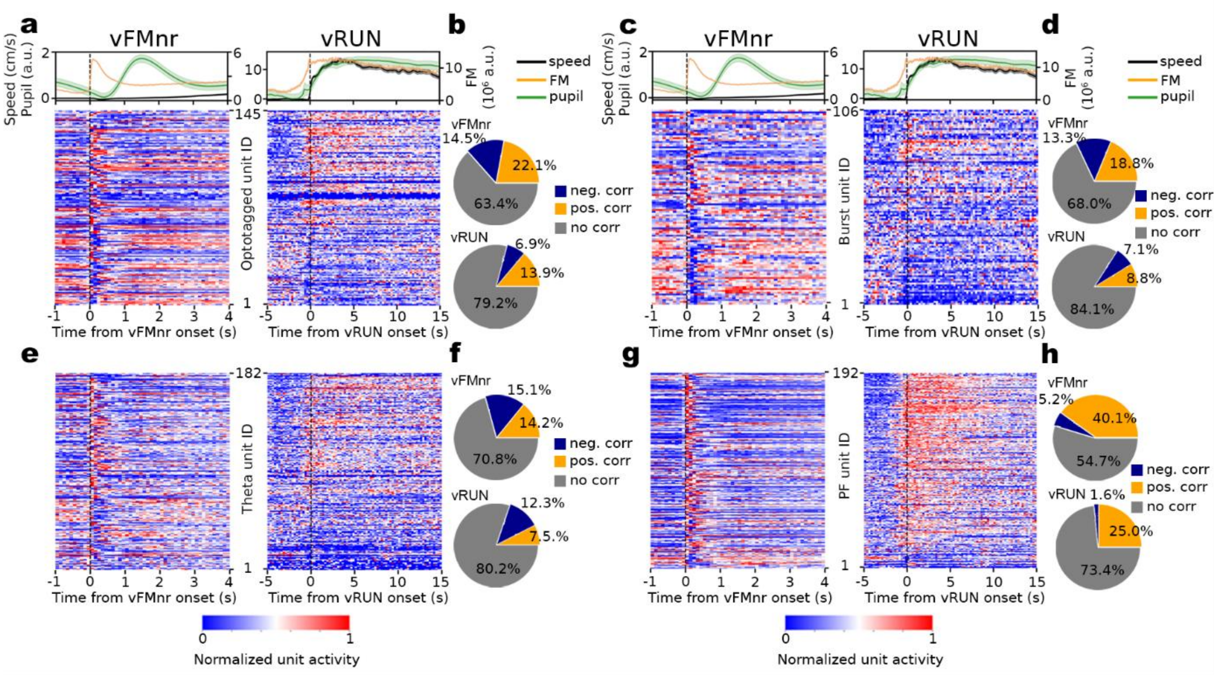


**Supplementary Fig. 3, Cell type specific modulation during voluntary FM and running. a** (top, left) Average speed, FM and pupil signals around voluntary stationary facial movement (vFMnr) onsets. (top, right) Average speed, FM and PUP signals around vRUN onsets, with (bottom) normalized activity of optotagged units sorted by modulation score. **b** Pie chart of positively and negatively correlated units of optotagged group with (top) vFMnr, and (bottom) vRUN. **c-d, e-f, g-h** Similar as in **a-b**, but with bursting, theta rhythmic, and persistent firing units, respectively.

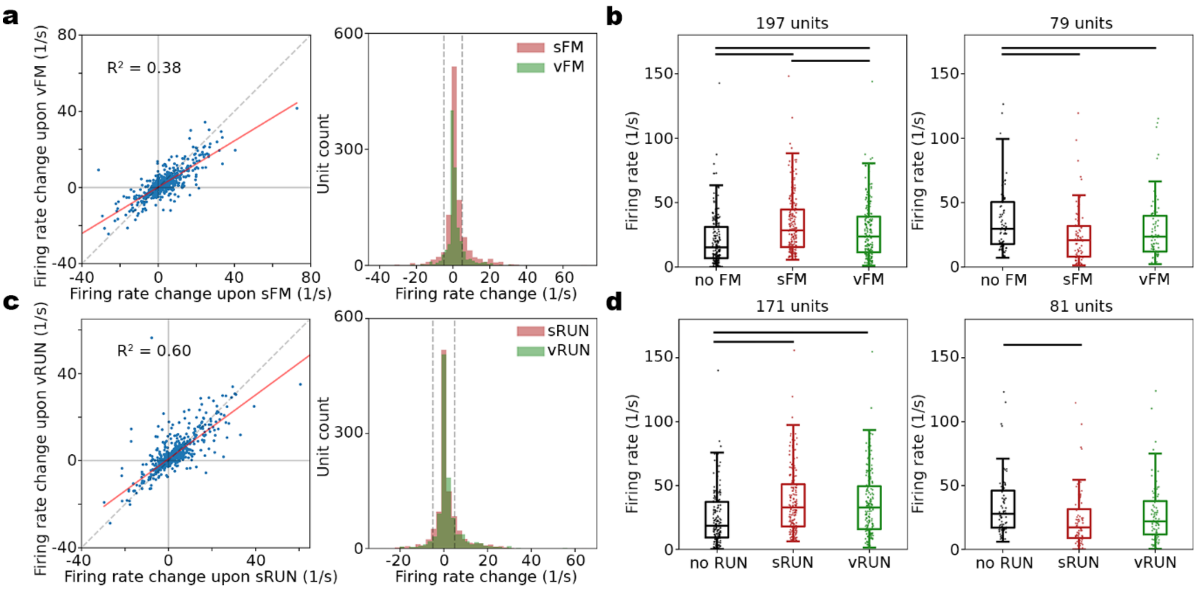


**Supplementary Figure 4, Recruitment of MSDB neuronal subgroups in voluntary and stimulated behaviors. a** (left) Firing rate change of recorded units from no FM to stimulated FM vs no FM to voluntary FM, (right) distribution of firing rate changes of units during voluntary compared to stimulated FMs, with (gray dashed lines) indication of 5 Hz firing rate threshold. **b** (left) Boxplot of firing rate of positively correlated units (over 5 Hz firing rate increase), and (right) boxplot of firing rate of negatively correlated units (over 5 Hz firing rate decrease). **c-d** Similar to **a-b**, but during sRUN and vRUN.

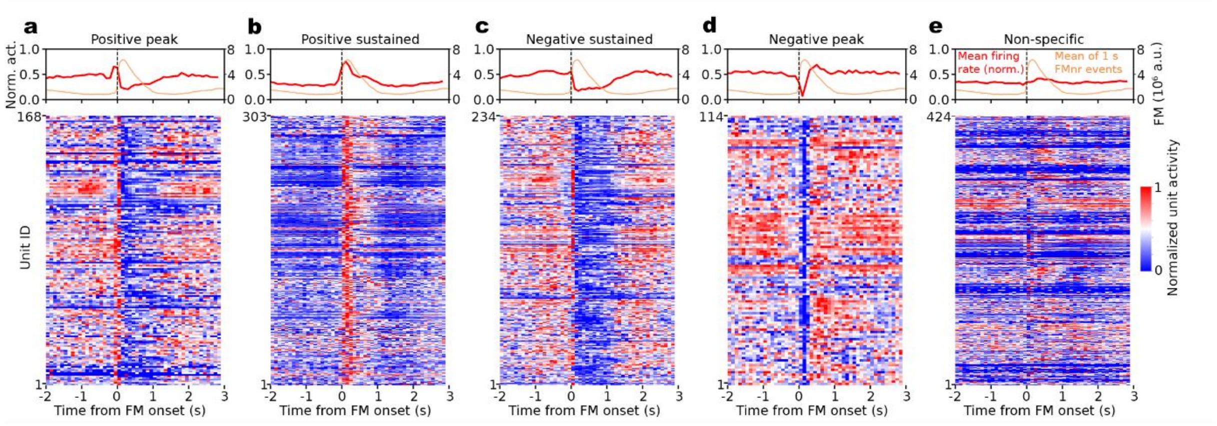


**Supplementary Figure 5, Distinct physiology of MSDB units during stationary facial movement. a-e** (top) Averaged (red) normalized unit activity, and (orange) 0.5-1s long FM through all experiments. (bottom) Heatmap of individual units in the presented activity phenotype groups.

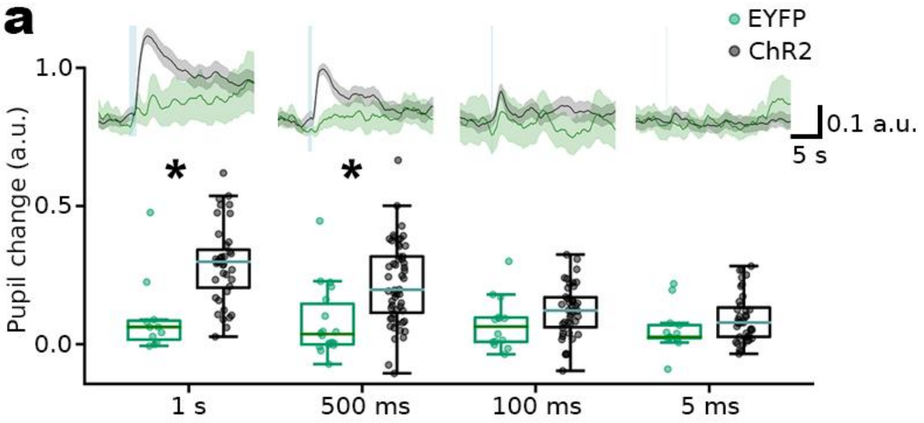


**Supplementary Figure 6, Pupil dilation is induced with shorter than 1s optical stimulation. a** Stimulation of MSDB VGluT2+ cells induce pupil dilation even with shorter stimulation durations. (top) Average pupil traces after various lengths of stimulation of ChR2 animals compared to EYFP control, (bottom) box plot of pupil change.
